# Reproducibility and Accuracy of Nanopore-Based Methylome Profiling of *Streptococcus dysgalactiae* subspecies *equisimilis* Strains from Cancer Patients

**DOI:** 10.1101/2025.06.04.657942

**Authors:** Taylor Schababerle, Omar Hayat, Jaeone Jung, Minh Le, Isabella Polic, Hanan Wees, Micah Bhatti, Samuel Shelburne, Xiaojun Liu, Awdhesh Kalia

## Abstract

**Background:** DNA methylation influences bacterial gene regulation, virulence, and restriction-modification (RM) systems. Advances by Oxford Nanopore Technologies (ONT) now enable direct methylome profiling from nanopore sequencing using the Dorado basecaller. However, the comparative performance of ONT-only versus hybrid-assembly reference-based methylation calling, particularly regarding genomic DNA quality and inter-operator variability, remains understudied.

**Methods:** Six operators independently prepared fifteen sequencing libraries each for nanopore (MinION R10.4.1 flow cells, Mk1D) and Illumina MiniSeq platforms for two *Streptococcus dysgalactiae subsp. equisimilis* strains (UT9728, 12 replicates; UT10237, 3 replicates). MicrobeMod v1.0.3 was used to identify methylation and motif profiles using Illumina-corrected hybrid reference assemblies (HRAs) and ONT-only reference assemblies (ORAs). Reproducibility and accuracy were compared using a custom genome annotation feature-enabled modular analysis that mapped and counted methylation site calls to CDS, rRNA and tRNA coordinates.

**Results:** Strain UT9728 predominantly exhibited N6-methyladenine (6mA) at GATC motifs, whereas strain UT10237 displayed dual methylation patterns: C5-methylcytosine (5mC) at CCWGG motifs and 6mA at GAGNNNNNTAA motifs. Both strains contained Type I and Type II RM systems; UT10237 uniquely harbored a Type IIG RM system with combined restriction and methylation activities. Motif identification concordance using HRAs and ORAs exceeded 99.9%. Reproducibility for methylation calls was high across independent replicates for both HRA (Pearson’s r >0.989) and ORA (Pearson’s r >0.993) methylation calls in GATC and CCWGG motifs but lower in the GAGNNNNNTAA motif (Pearson’s r (HRA) = 0.80; r (ORA) = 0.78). ORA-based methylation site calls for all motifs showed excellent precision and recall compared to HRA-based calls (F1-score >99.999%).

**Conclusion:** Our findings support the accuracy, robustness, and utility of ONT-only data based methylome profiling for bacterial epigenetic characterization. Our analytical framework facilitates detailed evaluations of reproducibility and accuracy.

**Data Summary and Availability:**

A. **Unprocessed_data_files**: DOI: <u>10.5281/zenodo.15555625</u>
  1. Raw paired-end FastQ Files (Illumina).
  2. Raw FastQ Files (ONT; unmodified basecalls).
  3. Raw uBAM Files (ONT; 6mA_5MC_modified_basecalls).
B. **Genome_assemblies_and_annotation_files**: DOI: <u>10.5281/zenodo.15558488</u>
  1. Genome_Assemblies reconstucted with ONT reads and polished with Illumina Reads (Hybrid assemblies);(strain_name_<rep_ID>_Hyb.gbk)
  2. Genome_Assemblies reconstructed with ONT reads and polished with ONT reads (ONT-only assemblies); (strain_name_<rep_ID>_ONT.fasta)
  3. GenBank flatfiles from Hybrid assemblies; (strain_name_<rep_ID>_Hyb.gbk)
  4. Genbank flatfiles from ONT-only assemblies. (strain_name_<rep_ID>_ONT.gbk)
C. **Data_Tables and Python_code** for modular analysis: DOI: doi.org/10.5281/zenodo.15579791
  1. 9728_10237_ORA+HRA_all_reps_microbemod_output.zip
  2. Methylation calls mapped to parsed genbank features & feature count matrix files.
    - 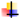 9728_HRA_reproducibility_analysis_tables.zip
    - 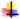 9728_ORA_reproducibility_analysis_tables.zip
    - 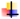 10237_HRA_5mC_6mA_reproducibility_analysis_tables.zip
    - 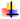 10237_ORA_reproducibility_analysis_tables.zip
  3. Python code for annotation feature-based modular analysis
    - 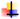 methylation_feature_hash_assert_FILTER_9728.py
    - 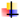 methylation_feature_hash_NOFILTER_9728.py
    - 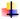 methylation_feature_hash_motif_10237.py
    - 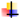 hyb_vs_ont_per_replicate_comparison.py
    - 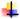 hyb_vs_ont_combined_plot.py
D. **Genome Assembly_Code Availibility**: https://github.com/TSchababerle/Bacterial-Methylation
  1. Bash script for automated Hybrid Genome Assembly
  2. Bash Script for automated ONT-Only Assembly

**Impact Statement:** This study provides a focused proof-of-principle demonstrating the robustness and reproducibility of Oxford Nanopore Technologies (ONT) sequencing-based methylome profiling without short-read correction. Using independent replicates prepared by multiple operators, we show that nanopore-only methods yield consistent, accurate methylation profiles concordant with short-read corrected ONT reads. This highlights the potential of ONT-only sequencing as a broadly accessible, reliable approach for rapid bacterial epigenomic characterization across diverse clinical contexts.

## Introduction

DNA methylation in bacteria, primarily involving N6-methyladenine (6mA) and C5-methylcytosine (5mC), is essential for gene regulation, host adaptation, and defense against foreign DNA via restriction-modification (RM) systems (1–3). RM systems, typically comprising methyltransferase (MTase) and restriction endonuclease (RE) enzymes, differentiate self-DNA by methylating specific motifs and cleaving unmethylated foreign DNA. Although RM genes usually occur in operons encoding both MTase and RE enzymes, orphan MTases lacking associated RE components exist and may have roles beyond DNA restriction defense, possibly acquired through horizontal gene transfer (4). Additionally, DNA methylation influences bacterial virulence by modulating genes linked to host invasion, antibiotic resistance, and uptake of mobile genetic elements (5–7). Thus, profiling methylation patterns in clinical isolates provides valuable insights into bacterial pathogenicity and genomic plasticity (8–10).

Traditional approaches, such as bisulfite conversion or Pacific Biosciences SMRT sequencing, have enabled base-resolution methylation mapping but can be labor-intensive or biased toward certain modifications. Recently, Oxford Nanopore Technologies (ONT) offers a relatively lower cost alternative by detecting unmodified and methylated DNA bases directly from native DNA. As DNA passes through a nanopore, methylated and unmodified bases cause characteristic disruptions in the ionic current, enabling detection of methylated bases without chemical conversion (11). Over the years ONT basecallers have steadily provided incrementally higher-quality sequence output enabling high quality genome reconstruction for genotyping bacterial strains without the need for error-correction by incorporating Illumina short-reads (12,13). Recent updates to ONT’s basecalling models, especially the Dorado software with integrated methylation-aware models, have substantially improved the ability to detect 6mA and 5mC with high confidence and resolution.

Despite these advances, studies leveraging the use of ONT-only workflows remain scarce, especially in clinical strains with complex RM systems or strain-specific epigenetic variation. While many studies still rely on hybrid approaches combining high-accuracy Illumina short reads with long-read assemblies for error correction, the increasing accuracy of ONT reads and tools like Dorado and MicrobeMod raise the possibility of complete genome and methylome assembly from nanopore data alone (14). Importantly, little is known about how strain-specific methylation profiles vary among closely related pathogens and how well ONT-only based detection captures this diversity in bacterial RM systems.

In this study, conducted as a part of the PaTHOGen-DS core curriculum in the graduate program in Diagnostic Genetics and Genomics at MD Anderson-School of Health Professions, we determine the reproducibility and accuracy of ONT only data for simultaneous genome and methylome reconstruction.

We used fifteen independent replicates from two clinical isolates of *Streptococcus dysgalactiae* subsp. *equisimilis* (SDSE), a β-hemolytic streptococcal species increasingly implicated in invasive infections, particularly in immunocompromised patients (15). We applied the Dorado v0.9.0 basecaller in combination with MicrobeMod v1.0.3 to identify and compare methylation motifs and RM system architectures between two SDSE strains isolated from cancer patients. Sequencing was performed using ONT MinION Mk1D R10.4.1 flow cells and an Illumina MiniSeq, enabling a direct comparison of Illumina-corrected hybrid reference assemblies (HRAs) and ONT-only reference assemblies (ORAs) as reference genome input for downstream methylome construction and characterization by MicrobeMod. Our findings reveal distinct methylation profiles between the two strains, including motifs associated with *Dam-* and *Dcm*-like activity and the presence of a Type IIG RM system unique to one isolate. We further demonstrate that ORAs, when generated with R10.4.1 chemistry and methylation-aware basecalling, may be sufficient for reproducible methylome reconstruction. By exploring strain-specific methylation in a clinically relevant pathogen and evaluating the performance of ORA- and HRA-based approaches, our work contributes to the growing toolkit for nanopore-based bacterial epigenomics.

## Materials and Methods

### A. Overall study design

To systematically evaluate how independent replication impacts genome assembly quality and methylome reconstruction in *Streptococcus dysgalactiae* subsp. *equisimilis* (SDSE), we conducted a two-phase experimental study using two clinical isolates (UT9728 and UT10237). In the first phase, eight biologically independent broth cultures were prepared by inoculating six single colonies of strain UT9728 and two single colonies of strain UT10237 into separate 5 mL aliquots of Todd-Hewitt broth, followed by overnight incubation. Genomic DNA (gDNA) was extracted from 1.5 mL of each broth culture, and samples meeting quality criteria (DNA Integrity Number [DIN] > 7) were sequenced using both Illumina and Oxford Nanopore Technologies (ONT). In the second phase, this experimental workflow was repeated with seven biological independent cultures (UT9728, six; UT10237, one), but DNA extraction was performed directly from colonies grown overnight on agar plates. This approach allowed us to assess whether the culture method influenced genome assembly and methylation analysis outcomes. ONT-only reference assemblies (ORAs) were constructed independently for each replicate and compared against hybrid reference assemblies (HRAs) generated separately for strains UT9728 and UT10237. This design enabled a controlled comparison of inter-replicate variation in assembly quality and methylome reconstruction under two commonly used sample preparation strategies.

### B. Bacterial strains and culture

SDSE isolates UT9728 (*serotype:G*; *emm-type*: stG166b.0; *mlst genotype*: ST360) and UT10237 (*serotype:C*; *emm-type*: stG62647.0; *mlst genotype*: ST20), originally isolated at MD Anderson clinical microbiology lab from cancer patients, were cultured on brain heart infusion (BHI) agar plates supplemented with 5% sheep blood (Hardy Diagnostics, Santa Maria, CA) for 18 hours at 37ºC. A single colony from each plate was then inoculated into 5mL of Todd-Hewitt broth (Hardy Diagnostics, Santa Maria, CA) and incubated for an additional 18 hours at 37ºC before gDNA extraction. To evaluate the influence of growth conditions on the quality of gDNA, genome assembly and methylome quality, 2-3 loopfuls of colonies grown overnight were resuspended in ice-cold phosphate-buffered saline (PBS, pH 7.0) and immediately processed for gDNA extraction.

### C. Genomic DNA (gDNA) extraction and quality control

gDNA was extracted from both broth- and plate-grown cultures using the GenElute™ Bacterial Genomic DNA Kit (Sigma-Aldrich, Cat. No. NA2120), supplemented with Mutanolysin (Sigma-Aldrich, Cat. No. M9901; stock concentration 5KU/mL) and Lysozyme (Millipore Sigma, Cat. No. L3790; stock concentration 10mg/mL) to enhance cell lysis. Manufacturer guidelines were followed, with vortexing minimized to prevent DNA shearing. DNA quality and concentration were assessed using the Qubit dsDNA BR Assay Kit (Life Technologies, Cat. No. Q32850) and TapeStation Genomic ScreenTape (Agilent, Cat. No. 5067-5365) along with corresponding reagents (Agilent, Cat. No. 5067-5366) (Table 1). In total, 15 gDNA samples were generated by six individuals: 12 from strain UT9728 and 3 from strain UT10237.

**Table 1:** Culture source, and gDNA and genome assembly metrics.

| Sample ID | Culture Source | Conc. (ng/μL) | DIN | HRA Genome Length | ORA Genome Length | HRA Assembly Coverage | ORA Assembly Coverage | SNV_1 | ORA Q-Score | SNV_2 |
| --- | --- | --- | --- | --- | --- | --- | --- | --- | --- | --- |
| UT9728_HW<sub>a</sub> | Plate | 78.4 | 9.1 | 2,092,990 | 2,128,112 | 128 | 449 | 15 | 51.5 | 23 |
| UT9238_HW<sub>b</sub> | Plate | 32.8 | 7.3 | 2,093,164 | 2,128,113 | 77 | 55 | 9 | 53.7 | 15 |
| U9728_IP<sub>a</sub> | Broth | 43.0 | 8.7 | 2,093,215 | 2,128,114 | 89 | 419 | 12 | 52.5 | 12 |
| UT9728_IP<sub>b</sub> | Broth | 70.2 | 8.5 | 2,093,166 | 2,128,113 | 85 | 133 | 11 | 52.9 | 11 |
| UT9728_IP<sub>c</sub> | Plate | 24.4 | 9.3 | 2,093,217 | 2,128,111 | 77 | 255 | 14 | 51.8 | 14 |
| UT9728_ML<sub>a</sub> | Broth | 39.6 | 9.2 | 2,092,990 | 2,128,113 | 84 | 214 | 14 | 51.8 | 13 |
| UT9728_ML<sub>b</sub> | Broth | 84.6 | 8.4 | 2,093,013 | 2,128,113 | 92 | 182 | 8 | 54.2 | 14 |
| UT9728_ML<sub>c</sub> | Plate | 22.4 | 8.2 | 2,093,164 | 2,128,112 | 71 | 290 | 13 | 52.1 | 17 |
| UT9728_OH<sub>a</sub> | Plate | 18.8 | 8.3 | 2,093,215 | 2,128,115 | 91 | 61 | 8 | 54.2 | REF |
| UT9728_OH<sub>b</sub> | Plate | 43.8 | 7.6 | 2,093,215 | 2,128,113 | 70 | 63 | 13 | 52.1 | 14 |
| UT9728_TS<sub>a</sub> | Broth | 19.3 | 8.5 | 2,093,215 | 2,128,114 | 112 | 161 | 12 | 52.5 | 18 |
| UT9728_TS<sub>b</sub> | Broth | 52.6 | 9.4 | 2,093,001 | 2,128,114 | 110 | 383 | 9 | 53.7 | 11 |
| UT10237_JJ<sub>a</sub> | Broth | 27.6 | 8.3 | 2,140,101 | 2,181,858 | 50 | 451 | 26 | 49.2 | REF |
| UT10237_JJ<sub>b</sub> | Broth | 34.0 | 8.9 | 2,139,990 | 2,181,857 | 65 | 200 | 24 | 49.6 | 26 |
| UT10237_TS<sub>c</sub> | Plate | 26.6 | 8.1 | 2,140,152 | 2,181,846 | 83 | 84 | 34 | 48.1 | 42 |

Conc. Qubit concentration (ng/μL), TapeStation DNA integrity number (DIN), and quality metrics for each of the replicates prepared in this study. \*SNV\_1: ONT-only assembly mapped with corresponding Illumina short-reads (ONT-only vs. Illumina reads); these SNVs were used to compute Q-scores. SNV\_2: ONT-only replicate assemblies compared to an arbitrarily chosen ONT-only assembly reference giving a measure of inter-replicate variance. REF= arbitrary reference.

### D. Whole genome Sequencing

All 15 DNA samples were sequenced using both Illumina short-read and Oxford Nanopore Technologies (ONT) long-read sequencing platforms. Short-read sequencing libraries were independently prepared by six individuals using the KAPA HyperPrep Kit (Roche, Cat. No. KK850), following the manufacturer’s protocol. Each library was quantified using the KAPA Library Quantification Kit (Roche, Cat. No. KK4835), pooled equimolarly based on measured concentrations, and sequenced on an Illumina MiniSeq. The sequencing run produced high-quality paired-end reads, achieving an average cluster density of approximately 185K/mm^2^, with > 90% of clusters passing quality filter. For long-read sequencing, libraries were individually prepared using the ONT Rapid Sequencing DNA v14 Barcoding Kit (ONT, Cat. No. SQK-RBK114.24) and sequenced on a MinION Mk1D equipped with a R10.4.1 flow cell. Libraries were also pooled equimolarly based on measured concentrations. The sequencing run generated approximately 17.34 Gb of basecalled data with an N50 of ~12.29kb. Basecalling of the raw nanopore data was performed using Dorado v0.9.0 software, in super accuracy mode (sampling rate: 5kHz), with the <u>dna_r10.4.1_e8.2_400bps_sup@v5.0.0</u> model on the Seadragon2 high-performance computing (HPC) cluster at MD Anderson, resulting in demultiplexed FASTQ files. For methylation analysis, additional basecalling was conducted using Dorado’s methylation-aware models: dna_r10.4.1_e8.2_400bps_sup@v5.0.0_4mC_5mC and dna_r10.4.1_e8.2_400bps_sup@v5.0.0_6mA. Methylation detection outputs were generated in BAM format for downstream methylome reconstruction and analysis. Raw sequence data including 1. Paired-end Illumina sequences (*_R1/R2.fastq.gz); 2. ONT basecalls-unmodified bases (*.fastq.gz) and 3. ONT basecalls – modified bases (*.bam) files are available at DOI: <u>10.5281/zenodo.15555625</u>.

### E. Genome Assembly, QC, and variation analysis

Hybrid reference assemblies (HRAs) and ONT-only reference assemblies (ORAs) were constructed for each of the 15 replicates. Illumina short-reads were quality-trimmed using fastp v.0.23.2 and combined with ONT long-read data using Unicycler v.0.23.2, followed by assembly polishing with Polypolish v.0.6.0 (16–18) to construct HRAs. Circularization and genome orientation steps were performed with Berokka v.0.2 and Dnaapler v.1.14.6, respectively, and gene annotation was conducted using Prokka v.1.14.6 (19–22). ORAs were generated using Flye v.2.9.1, followed by polishing with Medaka v.2.0.1 (23,24). Circularization, orientation correction, and annotation steps were identical to those used for HRAs. A bash script for automated implementation is available at (https://github.com/TSchababerle/Bacterial-Methylation). Genome assembly fasta files and PROKKA annotated Genbank files are available at DOI: doi.org/10.5281/zenodo.15558488. Pod5 files are available from authors upon request.

Coverage depth for hybrid genome assemblies was estimated during the Polypolish step and for ONT-only assemblies, during the Medaka polishing step. Assembly quality, including completeness and contiguity, were evaluated with QUAST v.5.2.0, BUSCO v.5.8.2 (25,26) and adaptor or foreign DNA contamination was assessed by NCBIs FCS-adaptor and FCS-GX toolkits, respectively (27) as implemented in Galaxy (28) (www.usegalaxy.org). Genetic variation between assemblies was assessed by pairwise alignment of all hybrid and ONT-only assemblies to arbitrarily chosen representative assemblies of each strain using Snippy v.4.6.0 to identify SNPs (29). The number of SNPs per assembly type were graphed for each strain on the software GraphPad Prism.

F. Methylation Calling

Methylation aware basecalling was performed using ONT’s Dorado basecaller which utilizes a neural network-based approach to differentiate modified from unmodified bases by analyzing ionic current disruptions during nanopore sequencing (11). Although Dorado supports detection of multiple DNA modifications including 4mC, 5mC, 6mA, 5mCG, and 5hmC, our analysis specifically focused on 5mC and 6mA because these are common modifications associated with bacterial RM systems. Original pod5 output files were reprocessed with methylation-aware models to generate methylation-specific unaligned BAM files. These BAM files were then converted to FASTQ format using Samtools v.0.3.3 and aligned against their respective assembled reference genomes using Minimap2 v.2.27 with parameters -T, MM and ML (30,31). We used two distinct sets of reference genome assemblies - HRAs and ORAs - as input for the MicrobeMod v.1.0.3 pipeline. Aligned BAM files were subsequently sorted and indexed with Samtools in preparation for downstream methylation analysis (30,32).

### G. Motif Identification

Assemblies aligned with methylation data were analyzed using MicrobeMod, a specialized computational tool providing two key outputs: (1) RM system gene annotation and (2) methylation-associated motif identification (32). RM gene annotation was conducted through the annotate_rm pipeline integrating Prodigal for gene prediction, HMMER and CATH-resolve-hits for protein domain searches, and sequence similarity searched using BLASTP against the REBASE database (33– 36).

Methylation motif discovery was performed using MicrobeMod’s call_methylation pipeline, which leverages Modkit for extracting methylation frequencies and the motif discovery algorithm STREME, a part of the MEME suite, for sensitive identification of sequence motifs (37,38). The Modkit module of MicrobeMod was run with default settings (66% methylation frequency with ≥ 10 supporting reads per call). Outputs from this pipeline included motif sequences, genome-wide methylation frequency statistics, and associate confidence scores. The final data outputs comprised three comprehensive tables, methylation site calls, motif occurrences, and RM system gene annotations, which are available at DOI: doi.org/10.5281/zenodo.15579790.

### H. Methylation Feature Mapping, Multi-Motif Benchmarking, and Comparative Analysis

A genome annotation derived feature-based analysis was preferred over all-site analysis because it provides biologically meaningful context thereby facilitating the interpretation and comparison of functional genomic regions across samples, and because nearly 95% of methylated sites were found in these feature-annotated regions.

Reproducibility and accuracy of methylation site calls across sequencing methods and replicates was evaluated using custom Python-based pipelines. Methylation site calls using ORAs and HRAs were mapped onto annotated genomic features (CDS, rRNA, tRNA) extracted from GenBank files. Feature classification distinguished named genes, hypothetical proteins (hashed by sequence identity), and RNA features. Strain with two methylation motifs (CCWGG for 5mC and GAGNNNNNTAA for 6mA) were analyzed separately.

Mapped methylation sites were summarized into per-feature counts, generating ‘all against all replicates’ feature count matrices (available at doi.org/10.5281/zenodo.15579791) and per-replicate feature-class summaries. These feature count matrices were then used to perform comparisons between sequencing platforms per-replicate using scatter and Bland-Altman plots to quantify concordance, bias, and agreement limits. Pearson correlation, Jaccard indices (presence-absence overlap), and Bland-Altman statistics quantified reproducibility. Additionally, confusion matrices were generated from feature-count matrices, calculating true positives, false positives, false negatives, and true negatives, along with sensitivity, specificity, precision, and F1 scores.

Feature-level reproducibility across replicates was assessed via Pearson correlation heatmaps and coefficient of variation (CV =SD/mean) normalizing dispersion across highly and sparsely methylated genes allowing quick identification of any outliers. UT9728 replicates were analyzed both with and without filtering for features detected in at least two replicates. Graphical outputs included per-replicate and across-replicate scatter plots, Bland-Altman plots, correlation heatmaps, and CV violin plots. All outputs and summaries were bundled into ZIP files for streamlined review.

Annotated Python scripts and MicrobeMod raw output, as well as all intermediate files from our analysis including, parsed gbk and MicrobeMod sites calls tables, presence/absence Jaccard matrices are available at <u>DOI 10.5281/zenodo.15579790; DOI 10.5281/zenodo.15558488</u>; DOI <u>10.5281/zenodo.15555625</u>.

## Results

### Replicate genome assemblies using ONT-only data consistently achieved ≥99.999% base call accuracy

Despite differences in culture source, DNA input concentration, DNA integrity numbers (DIN 7.3-9.4), and multiple operators, high-quality scores (Q48+) and 99.2% BUSCO completeness (Figure S1) were consistently obtained across all genome assemblies (Table 1). No adaptor or foreign DNA contamination was detected. Reflecting these high Q-scores, only a modest number of errors (SNVs/Indels) were identified in replicate ORAs using Illumina short-reads: 9–16 SNVs in UT9728 replicates and 24–34 SNVs in UT10237. Inter-replicate comparisons among ORAs also showed variability despite identical processing conditions (Table 1, SNP2). In contrast, HRAs from both strains exhibited no SNVs or indels, underscoring their high technical reproducibility independent of all differences noted above (Figure S2a, S2b). Moreover, most SNVs/Indels clustered within ± 10 bp of methylated sites, consistent with previous observations (Figure 2c). However, ORAs demonstrated remarkable consistency in genome size (within 3–12 bp), while HRAs varied slightly more (range: 111–227 bp), averaging approximately 35–40 kb shorter, likely due to differential resolution of repeat or rRNA regions. Thus, we conclude that ORAs can reliably produce structurally complete genomes of consistent sizes irrespective of input DNA variability, integrity or culture method for the two SDSE strains in included in this study.

**Figure 1:**
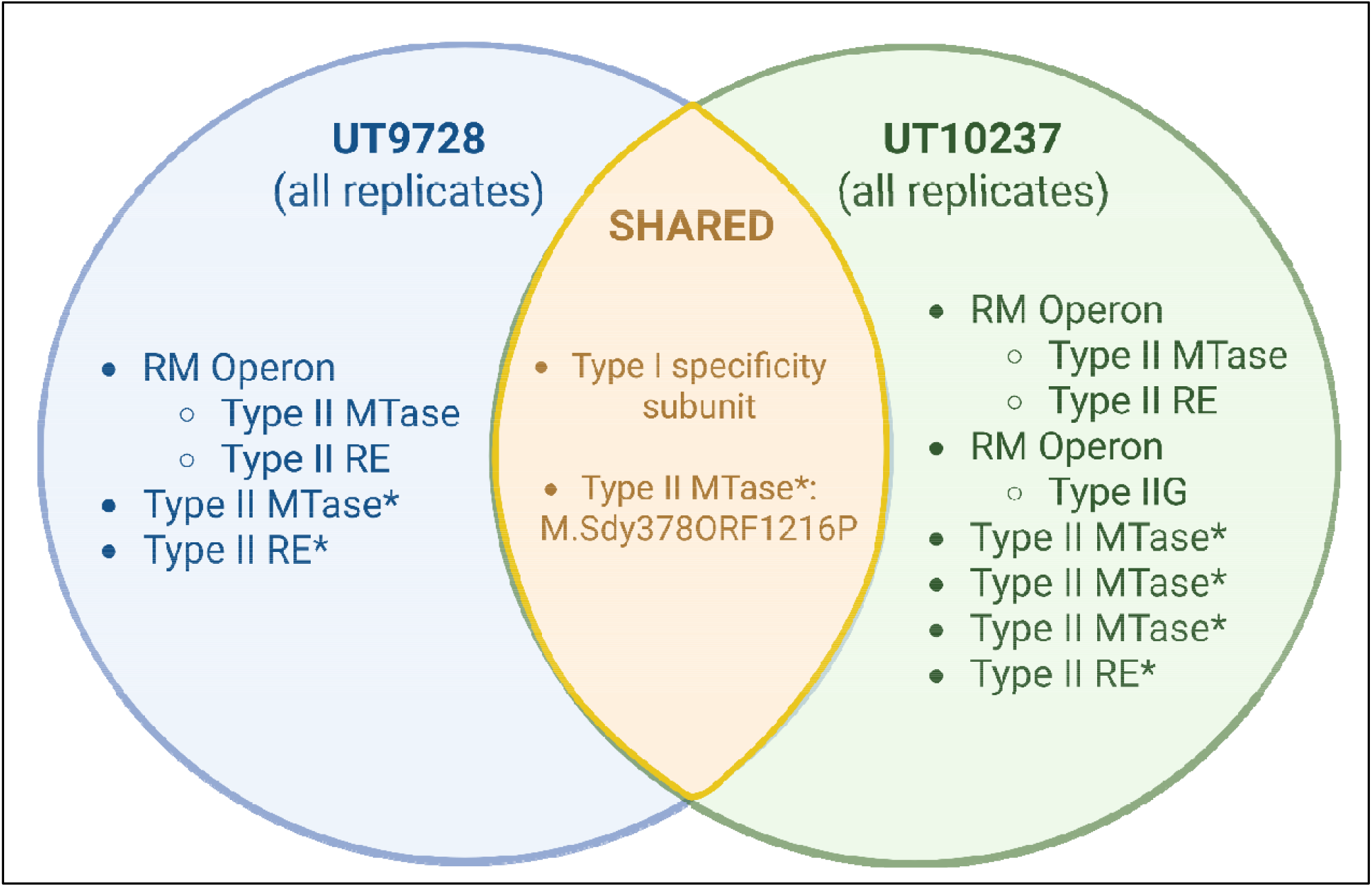
Shared and unique RM systems identified in the two SDSE strains. Venn diagram detailing the RM systems for strains UT9728 and UT10237 and illustrating the common MTase and Type I specificity subunit. * Denotes an orphan MTase or RE.

**Figure 2:**
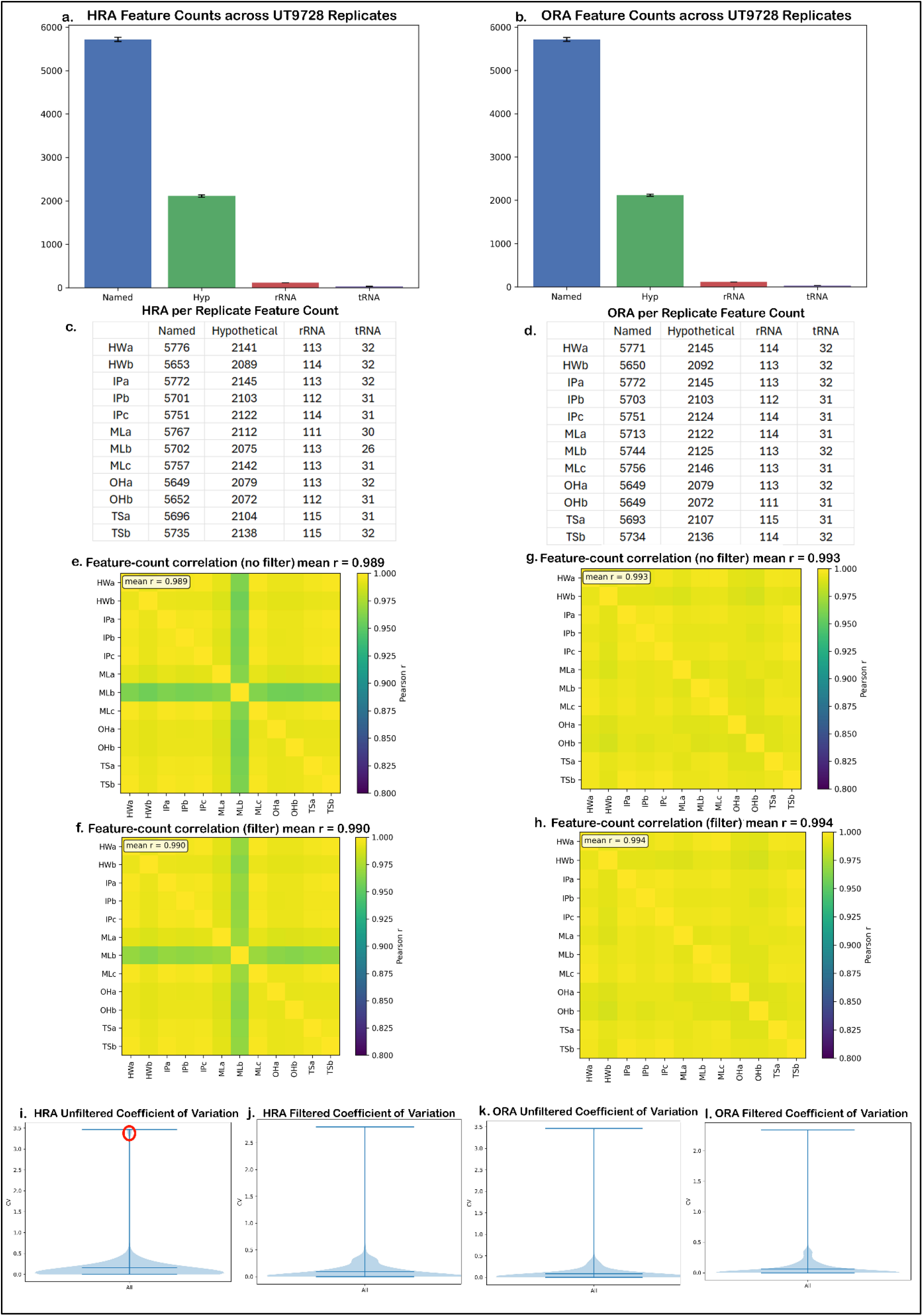
High reproducibility of feature-level methylation profiles across twelve SDSE UT9728 ONT-only replicates. (a–d) Distribution of total methylation site counts mapped to annotated genomic features using the hybrid reference assembly (HRA; panels a, c) or ONT-only reference assembly (ORA; panels b, d). Panels (c) and (d) show per-replicate distributions, highlighting consistent feature-level methylation across all 12 assemblies. (e–h) Pearson correlation heatmaps of all 66 pairwise comparisons of feature-level methylation counts across replicates. Panels (e) and (f) show HRA-based comparisons before and after singleton site filtering; panels (g) and (h) show the same for ORA-based calls. Removal of singleton calls improves replicate concordance, particularly in low-frequency features. (i–l) Violin plots displaying the coefficient of variation (CV) in methylation site counts per feature across replicates. Panels (i) and (j) show HRA-based values before and after singleton filtering; (k) and (l) depict ORA-based results. Red circle in panel (i) denotes an outlier replicate with elevated dispersion, despite high Q-score. Overall, low CV values and strong inter-replicate correlations (Pearson r ≥ 0.989) confirm the technical reproducibility of feature-level methylation calls across assemblies.

### Strain-Specific Methylation Signatures and Distinct RM System Architectures in SDSE Genome

We next evaluated whether ORAs could accurately capture and distinguish the methylation profiles of two SDSE strains. In all replicate HRAs and ORAs, MicrobeMod consistently identified only 6mA methylation at GATC motifs in strain UT9728, indicative of Dam-like methyltransferase activity (Figure S3a). In contrast, strain UT10237 exhibited a more complex methylation landscape, with 5mC at CCWGG motifs consistent with Dcm-like activity (Figure S3b), and 6mA at GAGNNNNNTAA degenerate motifs associated with a Type IIG RM system (Figure S3c).

Methylation motif calls made using MicrobeMod showed high concordance between HRA and ORA datasets. For strain UT9728, concordance values for the GATC motif ranged from 99.8% to 99.9% (Table S1). For UT10237, values ranged from 99.9% to 101.6% for GAGNNNNNTAA-suggesting slight overcalling with ORAs- and 99.1% to 100% for CCWGG (Table S1). While minor variation in motif counts was observed among ORA replicates, the proportion of modified sites remained consistent (Table S2b) reinforcing that basecalling, not mapping, dominates quantitative accuracy. Furthermore, strand bias metrics were flat across all conditions (Tables S2a and S2b), arguing that Dorado’s homopolymer improvements have largely solved the forward-reverse imbalance that plagued earlier R9 chemistry.

Collectively, these distinct and reproducible methylation signatures indicate ORAs when used with MicrobeMod can recover biologically meaningful methylome variation in SDSE genomes. Overall, results underscore the robustness of MicrobeMod’s motif detection across sequencing strategies.

MicrobeMod analysis identified six RM genes in SDSE strain UT9728, three of which matched known homologs in the REBASE database (Figure 1). Two of these formed a Type II RM operon comprising a methyltransferase (MTase; M.Sdy89ORF13680P), and a restriction endonuclease (RE; Pst19aORF1373P), showing high sequence identities - 93.66% (E-value of 5.4E-104) and 96.06% (E-value 1.1E-79), respectively - strongly supporting their functional annotation. This operon recognizes the GATC motif in the genome, consistent with our methylation analyses. Two additional Type II orphan MTases were identified, including one with 100% identity to the REBASE-listed M.Sdy378ORF1216P (E-value 4.3E-36) suggesting that it is likely SDSE-specific. A solitary Type II RE lacking a known homolog was also identified, possibly an inactive remnant or missing its cognate MTase. A Type I specificity subunit was identfied without accompanying Type I system components, suggesting gene loss or recent horizontal gene transfer.

Strain UT10237 harbored nine RM genes: two operons and six singletons (Figure 1). The first operon encodes a Type II RM system with MTase (M.Sdy7136ORF314P) and RE (Sdy7136ORF314P), both with 100% identity to SDSE homologs (E-values 2.7E-82 and 7.6E-21). The second encodes a Type IIG RM system-a single polypeptide with both MTase and RE activity-identical to Sdy7136I and targeting the GAGNNNNNTAA motif seen in methylation data. Four Type II orphan MTases were identified, two matching known homologs (M.Sps20026ORF1737P and M.Sdy378ORF1216P). Like UT9728, UT10237 also contains a singleton RE gene and a standalone Type I specificity subunit. These distinct but partially overlapping RM repertoires may highlight strain-specific ecology and defense strategies with implications for methylation patterns, genome stability, and pathogenicity.

### High reproducibility of HRA- and ORA-based methylation site counts across UT9728 replicates

Because strain UT9728 contains a single GATC motif and many replicates, we first assessed the reproducibility of MicrobeMod methylation calls within each assembly type. Methylation sites were mapped to GenBank-annotated genomic features—named and hypothetical coding sequences (CDS), rRNAs, and tRNAs—and site counts were summed per feature per replicate (Tables S2a and S2b). Counts were highly reproducible across all 12 ORA replicates (Figure 2). Both HRA and ORA datasets displayed similar feature-type distributions, with named genes harboring the most sites, followed by hypothetical CDS, rRNAs and tRNAs (Figures 2a, 2b). Nevertheless, feature-level counts varied modestly among replicates (Figures 2c, 2d).

To quantify consistency, we generated a feature-by-replicate count matrix and performed pairwise comparisons. Pearson correlations exceeded 0.98 for all 66 pairwise comparisons in both HRA and ORA datasets (Figures 2e–2h), although the HRA MLb replicate showed a noticeable drop (Figure 2e). Coefficient-of-variation (CV) analysis revealed that most features were tightly distributed (CV < 0.3), with a few high-CV outliers (~3.5; Figure 2i). Removing singleton methylation calls (sites present in only one replicate) raised the mean HRA correlation from 0.98 to 0.99 (Figure 2f) and eliminated the high-CV outliers (Figure 2j). ORA replicates displayed even higher correlations both before and after filtering (Figures 2g, 2h; 2k, 2l). Collectively, these results demonstrate robust, reproducible methylation profiling within each assembly type, supporting the reliability of the feature-based approach.

### Strong concordance between ORA- and HRA-based methylation site calls

Concordance between ORA- and HRA-based methylation site calls were evaluated across all UT9728 replicates. Feature-level site counts showed excellent agreement between assembly methods (Figure 3). Benchmarking with Pearson correlation, Jaccard index and Bland-Altman analysis confirmed this concordance (Figure 3a). Replicate-specific Pearson coefficients ranged from 0.964 (MLb) to 1.000 (IPb, OHa), indicating near-perfect linear relationships. Scatter plots of pooled (Figure 3b) and individual replicates (Figure S4) further highlight the close alignment of counts.

**Figure 3:**
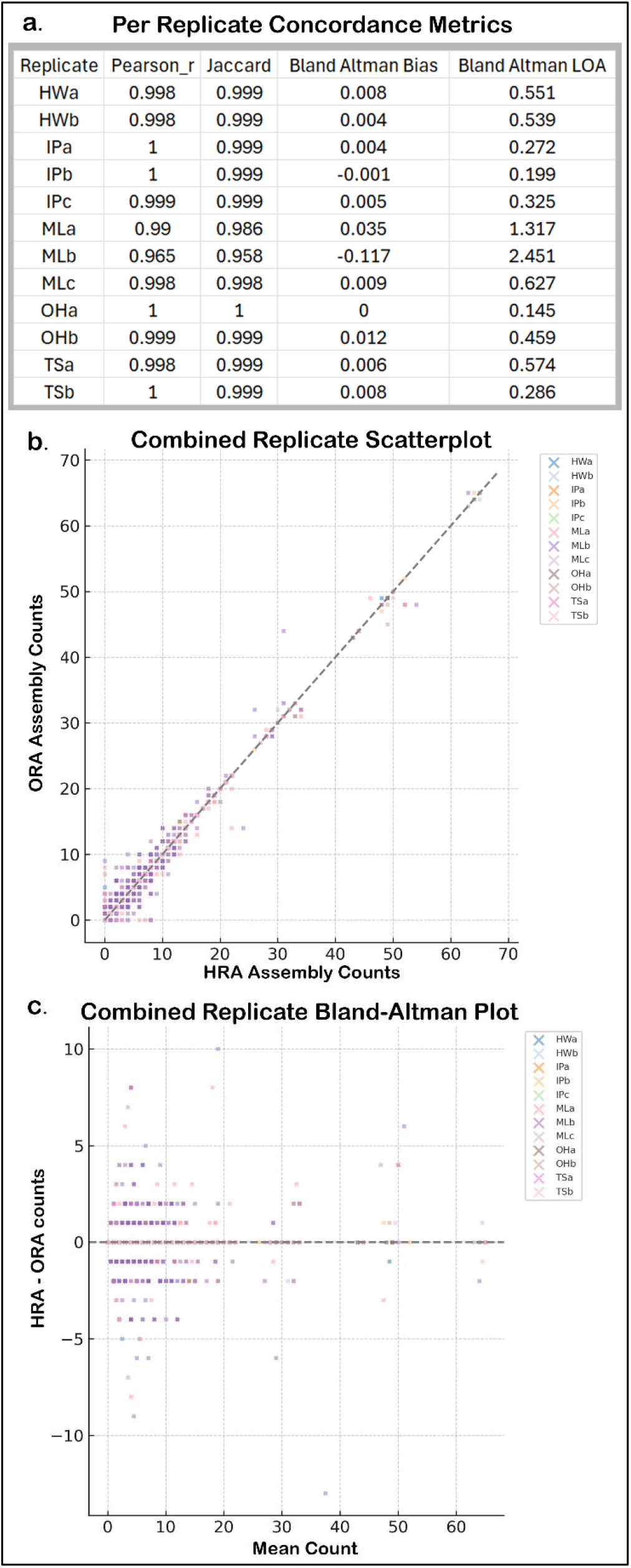
High concordance between ORA- and HRA-based methylation calls in SDSE strain UT9728. (a) Summary table showing global agreement between feature-level methylation site counts derived from ORAs and HRAs, assessed using three metrics: Pearson correlation coefficient (linear association), Jaccard index (feature overlap), and Bland–Altman bias with 95% limits of agreement (LOA). (b) Scatterplot comparing per-feature methylation site counts between ORA and HRA across all 12 replicate pairs. Most points fall tightly along the identity line, indicating strong concordance; the observed dispersion is primarily driven by a single outlier replicate (MLb), with some contribution from MLa. (c) Bland–Altman plot showing the difference in methylation counts (HRA-ORA) versus their mean per feature, aggregated across all replicates. The average bias is low, but LOA exceeds ±1, largely due to replicate MLb and, to a lesser extent, MLa. Individual replicate plots are available in Supplementary Figures S4 and S5.

Bland-Altman analysis revealed minimal bias and narrow limits of agreement for most replicates (Figure 3c). This was especially exemplified by HWa (mean difference ≈ 0, limits ±1.96 SD = 0.65; Figure S5a). Similar low-bias, tight-limit patterns were observed in other replicates. The sole outlier was MLb, which showed wider limits (±1.96 SD = 2.49) and a modest negative bias (−0.11), indicating greater dispersion between ORA and HRA counts (Figure S5e). Outliers predominantly occurred in features with low mean site counts, where small absolute differences inflate proportional discrepancies but have little effect on global metrics.

HRA- and ORA-based methylation profiles showed strong concordance: we detected few false positives, false negatives, and true negatives - defined here as features not methylated in the paired HRA-ORA comparison but methylated in other replicates (Table S3a). Thus, ORA call achieved high recall (range 0.96 – 1) and accuracy (0.98 – 1). Taken together, ORA-based feature-specific methylation site counts faithfully recapitulate HRA-based methylation profiles, with high concordance across replicates and only isolated variability attributable to technical factors in specific datasets.

### Reproducibility and accuracy of motif-specific methylation calls in strain UT10237

Strain UT10237 contains two distinct motifs -CCWGG (5mC) and the longer, degenerate GAGNNNNNTAA (6mA) - allowing assessment of how motif complexity influences the consistency of methylation profiling. HRA- and ORA-based MicrobeMod calls were mapped to GenBank-annotated features, and site counts were summed *per motif per feature per replicate* - three replicates per assembly type. (Tables S2a and S2b).

#### CCWGG motif (5mC)

Feature-type distributions were nearly identical between HRA (Figure 4a) and ORA datasets (Figure 4b). Within each assembly type, per-replicate profiles varied slightly (Figures 4e and 4f). Pairwise replicate comparisons confirmed this reproducibility with HRA-based calls displaying a Pearson’s mean r = 0.98; and ORA-based calls displaying a Pearson’s mean r = 0.97 (Figures 4i and 4j).

**Figure 4:**
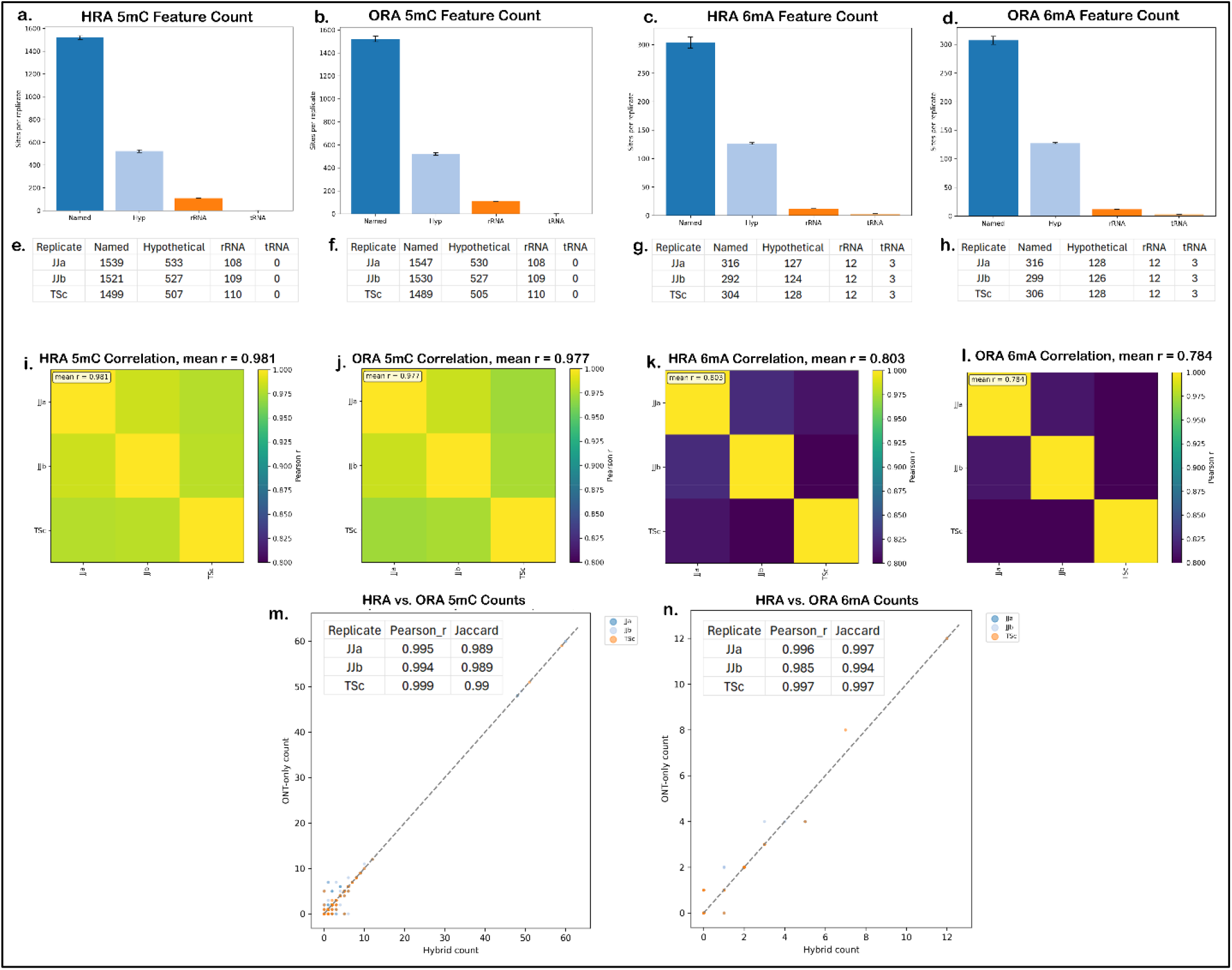
Reproducibility of methylation counts in two distinct motifs within SSDSE strain UT10237: the stable CCWGG (5mC) motif and the more degenerate GAGNNNNNTAA (6mA) motif. (a–d) Distribution of total methylation site counts mapped to annotated genomic features, comparing HRA and ORA. Panels (a, c) represent HRA-based distributions; (b, d) show corresponding ORA-based distributions. (e–h) Per-replicate distributions of feature-level methylation counts, illustrating consistency across replicates. Panels (e, g) display HRA-based replicate distributions; (f, h) depict ORA-based distributions. Greater consistency is evident for the CCWGG motif relative to the degenerate GAGNNNNNTAA motif. (i–l) Pearson correlation heatmaps summarizing all 6 pairwise comparisons of feature-level methylation site counts across the three available replicates. Correlations are shown separately for the stable CCWGG motif (panels i, j) and the degenerate GAGNNNNNTAA motif (panels k, l), and for both HRA-based (i, k) and ORA-based (j, l) calls. Stronger replicate correlations are evident for CCWGG motif across both assembly methods. (m–n) Scatterplots comparing per-feature methylation site counts between ORA and HRA across replicates, summarized by Pearson’s correlation coefficient (r) and Jaccard index (insets). Panel (m) shows high concordance for the CCWGG motif, while panel (n) illustrates lower reproducibility and greater scatter in the degenerate GAGNNNNNTAA motif.

#### GAGNNNNNTAA motif (6mA)

By contrast, the degenerate 6 mA motif exhibited greater variability. Although overall feature distributions remained similar between HRA- and ORA-based calls (Figures 4c, 4d, 4g and 4h), per-replicate patterns diverged more noticeably. Pearson correlations dropped to 0.80 in HRA-based calls and 0.78 in ORA-based calls (Figures 4k and 4l), indicating reduced concordance relative to the CCWGG motif. Variability was most pronounced in features with low mean site counts (Tables S3b and S3c), suggesting stochastic detection of sparse GAGNNNNNTAA sites.

Because only three replicates were available, coefficient-of-variation and Bland-Altman analyses were not performed for UT10237. Nonetheless, the correlation data show that motif composition markedly affects reproducibility: the simple, densely represented CCWGG motif was called with high precision (Pearson’s mean r, range = 0.994 – 0.999), whereas the longer, degenerate GAGNNNNNTAA motif showed greater variability across replicates (Pearson’s mean r, range = 0.985 – 0.997) (Figures 4m and 4n). Even so, ORA-based calls accurately recapitulate HRA-based patterns for both motifs with high precision and recall (Tables S3b and S3c), supporting their use for comprehensive methylome profiling in genomes containing multiple, structurally diverse motifs.

## Discussion

Many recent studies have demonstrated the utility (12,13) and limitations (39) of ONT-only genome assemblies from various bacterial species. However, few studies have evaluated nanopore-sequencing for methylome profiling with the newer advanced Dorado models and R10 flow-cell chemistry (14,40,41). Our analysis aimed to assess the feasibility and reproducibility of using ORAs as reference during MicrobeMod analysis for characterizing the methylome of two clinical strains of SDSE, UT9728 and UT10237. Using fifteen independent ONT and Illumina datasets from two *Streptococcus dysgalactiae* strains, we show that Dorado-based ONT-only assemblies (ORAs) deliver nearly identical methylome profiles to Illumina-polished hybrids (HRAs). UT9728 carries a canonical *Dam*-like 6mA system targeting GATC sites, whereas UT10237 combines a *Dcm*-like 5mC system at CCWGG with a sparse, degenerate 6mA motif (GAGNNNNNTAA) encoded by a Type IIG RM enzyme. Feature-level methylation counts were highly reproducible across replicates (Pearson r > 0.98 for simple motifs) and ORA/HRA calls were almost perfectly concordant (F1 > 0.999). The only notable variance was confined to one hybrid replicate (MLb) and to the long degenerate motif, highlighting how motif complexity and technical bias, rather than read accuracy, now limit ONT methylome studies.

Figure 1 underscores two contrasting epigenomic strategies. UT9728 encodes a compact Type II GATC RM pair plus orphan *Dam*-like MTases, yielding dense 6mA methylation marks across coding genes, ~110 rRNA sites and 32 tRNAs sites. *Dam* methylation synchronizes replication, directs MutHLS mismatch repair, and toggles virulence genes, giving rapidly growing strains a flexible “transcriptional clutch” (42). In UT10237, a *Dcm*-like system methylates ~110 rRNA sites but no tRNAs, consistent with a steadier metabolic style in which 5mC primarily serves genome defense and fine-scale expression tuning (43). The additional Type IIG enzyme creates the 22-bp GAGNNNNNTAA motif; its low occupancy suggests phase-variable expression or decay, mirroring patterns seen in recent methylome surveys that link IIG systems to niche-specific phage pressure (14). Moreover, both strains harbored a strain-specific repertoire of orphan type II MTases, which usually do not cleave unmethylated foreign DNA but can mediate crucial regulatory roles by modulating bacterial virulence gene expression or other physiological processes, e.g., mobile element activity (44). We speculate that strain-specific differences indicate how exchanging Dam for Dcm and IIG could influence both defense mechanisms and regulatory potential. UT9728 appears to maintain permeability to horizontal gene transfer and may be poised for rapid growth, whereas UT10237 seems to employ a more stringent RM barrier while potentially retaining an auxiliary 6mA ‘bet-hedge’ switch. Further experiments will be needed to test these hypotheses and refine our understanding of functional implications.

### Reproducibility of simple motifs

In UT9728, feature-level GATC-6mA counts were remarkably consistent (Pearson r ≥ 0.989 across all 66 pairwise comparisons), and ORA-versus-HRA scatterplots lay tightly on the identity line (Figures 2, 3). This performance matches the cross-laboratory work of Galeone et al., who resequenced the same isolates in three institutes and showed that motif recall rises steeply with depth but plateaus once genome-wide coverage exceeds ≈50 ×, with only marginal gains beyond 100 × (14). Because every one of our 24 assemblies sits well above that plateau, near-perfect correlations are expected, confirming that coverage - not operator - is the dominant factor in reproducibility.

Although the MLb hybrid replicate assembly exhibits high base call accuracy (Q-score 54, ~99.999%), the elevated outliers and CV distribution indicate the presence of systematic bias rather than random error. This suggests that consistent technical factors - such as systematic errors in methylation calling algorithms - may be influencing the detection of methylation sites in this replicate. Such biases could lead to reproducible deviations affecting specific genomic features or sequence contexts, resulting in the observed long tail in CV values. These findings highlight that even with excellent base-level accuracy, systematic technical biases can impact methylation site reproducibility and emphasize the importance of addressing these factors through improved experimental design and computational normalization to ensure robust methylation profiling.

### Complex, degenerate motifs remain challenging

For UT10237, feature-level concordance for the compact CCWGG-5mC motif (r ≈ 0.98) paralleled UT9728, yet the degenerate GAGNNNNNTAA-6mA motif fell to r ≈ 0.80 across replicates despite near-identical ORA-vs-HRA means. The MicrobeMod developers previously reported that degenerate motifs overlapping CCWGG produce unstable STREME scores, and variable locus counts when motif instances are sparse (32). Our data confirm that observation in a clinical isolate. Because the motif is targeted by a single Type IIG polypeptide whose expression may itself phase-shift, a biological contribution to variance cannot be excluded; however, the stable ORA/HRA agreement (F1 ≈ 0.998) indicates that algorithmic sensitivity, not platform accuracy, is the present bottleneck.

Our study has a few limitation**s:**

#### Taxonomic scope

Our two-strain design was sufficient to benchmark pipeline reproducibility but cannot capture species-wide variation in RM architecture or motif composition in SDSE. Galeone et al. advocate large, phylogenetically diverse panels to disentangle technology-specific artefacts from lineage-specific biology (14); we concur and view our dataset as a focused proof-of-principle.

### Assembly strategy

We deliberately used a single, widely adopted genome reconstruction route (Flye + Medaka or Unicycler + Polypolish). The observed variation in ONT-only assemblies likely stems from using Medaka 2.0.1 without the --bacteria option, which enables methylation-aware polishing. Since this study focuses on methylation rather than genotyping, we reason that omitting this option should not significantly impact per-replicate concordance or correlations between ORA- and HRA-based methylation calls. However, because basecalling accuracy may influence motif/site calling—particularly for long degenerate motifs—we acknowledge this as a potential limitation, given that methylation site call correlation in ONT assemblies was slightly lower than in hybrid assemblies (0.78 vs 0.80) for strain UT10237.

### Feature scope

By constraining analysis to coding sequences, rRNAs, and tRNAs (~95 % of calls), we avoided alignment ambiguities in low-complexity intergenic DNA. Yet adaptive methylation of promoter islands or mobile elements will be missed. Pilot surveys in *E. coli* and *Listeria* show that intergenic 6mA contributes disproportionately to phase variation; adding these regions could either dilute correlation (if noisier) or tighten it (if systematically modified). Incorporating calibrated confidence thresholds for low-depth intergenic sites, as suggested by MicrobeMod, will be essential.

In conclusion, our study provides a focused proof of principle for the robustness and reproducibility of using ONT reads alone—without short-read error correction—by achieving highly similar results across replicates prepared independently by six individuals. Despite differences in operator techniques and expertise, the outcomes remained consistent, highlighting that technical reproducibility is not dependent on a single individual performing the tasks.

## Author Contributions

TS and AK - conceptualization, data curation, formal analysis, methodology, software, validation, visualization, writing–original draft, and writing–review and editing. TS, HW, IP, ML, OH, and JJ – generated sequencing data and to writing review and editing. XL and AK – methodology training, project administration, resources, supervision, and writing–review and editing. MB and SAS - provided resources and participated in writing–review and editing.

## Acknowledgements

We thank the Peter and Cythia Hu Cardinal Health Scholarship (TS and JJ); the Kinder Foundation Endowed Scholarship (IP); the McKowen Scholarship (ML); the Dell Scholarship (OH); and the Fulbright Foundation (HW) for funding support. The authors acknowledge the support of the High-Performance Computing for research facility at the University of Texas MD Anderson Cancer Center for providing computational resources that have contributed to the research results reported in this paper. Figure 1 was created using BioRender (www.biorender.com).

