## Supplementary Figures S1-S5 & Tables S1-S3 for "Reproducibility and Accuracy of Nanopore-Based Methylome Profiling of *Streptococcus dysgalactiae* subspecies *equisimilis* Strains from Cancer Patients"

**2, Department of Lab Medicine, The University of Texas MD Anderson Cancer Center**

**3, Departments of Infectious Diseases and Genomic Medicine, The University of Texas MD Anderson Cancer Center**

**Author for Correspondence:**

Awdhesh Kalia, PhD

**List of Supplementary Figures**

Figure S1: BUSCO

Figure S2: HRA vs ORA SNVs

Figure S3: IGV Snapshot

Figure S4: UT 9728 individual scatter plots

Figure S5: UT9728 individual BA plots

**List of Supplementary Tables**

Table S1: Motif Call Concordance and Percent Modified Sites

Table S2: Total Sites and Strand Bias Table

Table S3: Confusion Matrix from MSC Analysis

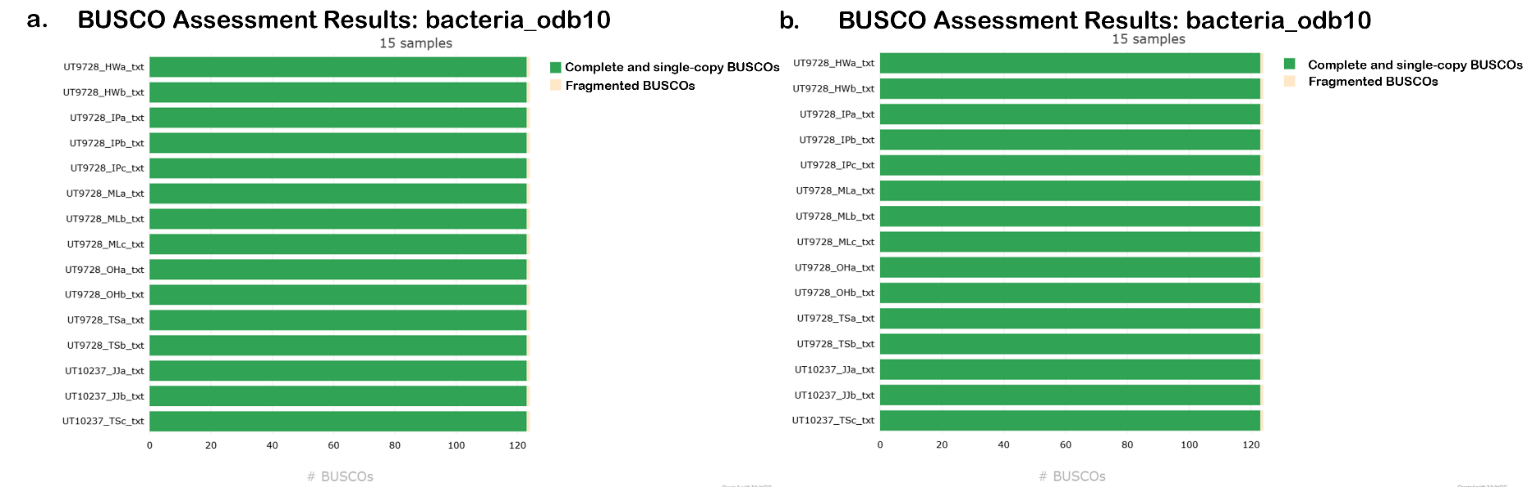
**Supplementary Figures**

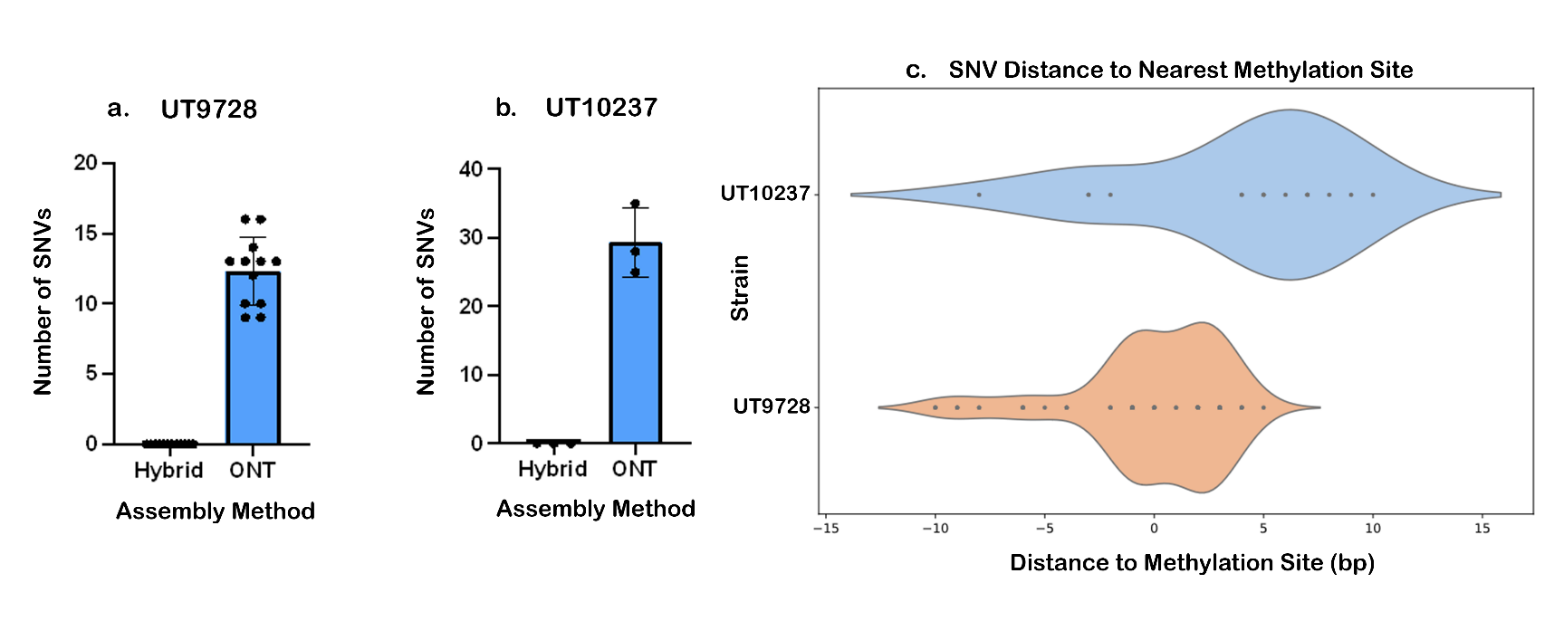
**Figure S1: BUSCO assessment results for hybrid and ONT-only assemblies.** (a) Hybrid assembly BUSCO assessments for all 15 replicates were 99.2% complete. (b) ONT-only assembly BUSCO assessments for all 15 replicates were 99.2% complete.

**Figure S2. Independent** **ONT-only assembly replicates show variable genetic differences.** (a) Bar graph showing the SNV counts between hybrid and ONT-only assemblies for strain UT9728. (b) Bar graph showing the SNV counts between hybrid and ONT-only assemblies for strain UT10237. (c) Violin plot showing the localization of SNVs to methylation sites in both strains. SNV sites were manually assessed for colocalization with methylation sites within ±10 base pairs. All replicates with identified sites were aggregated into their respective strain and graphed using the seaborn library.

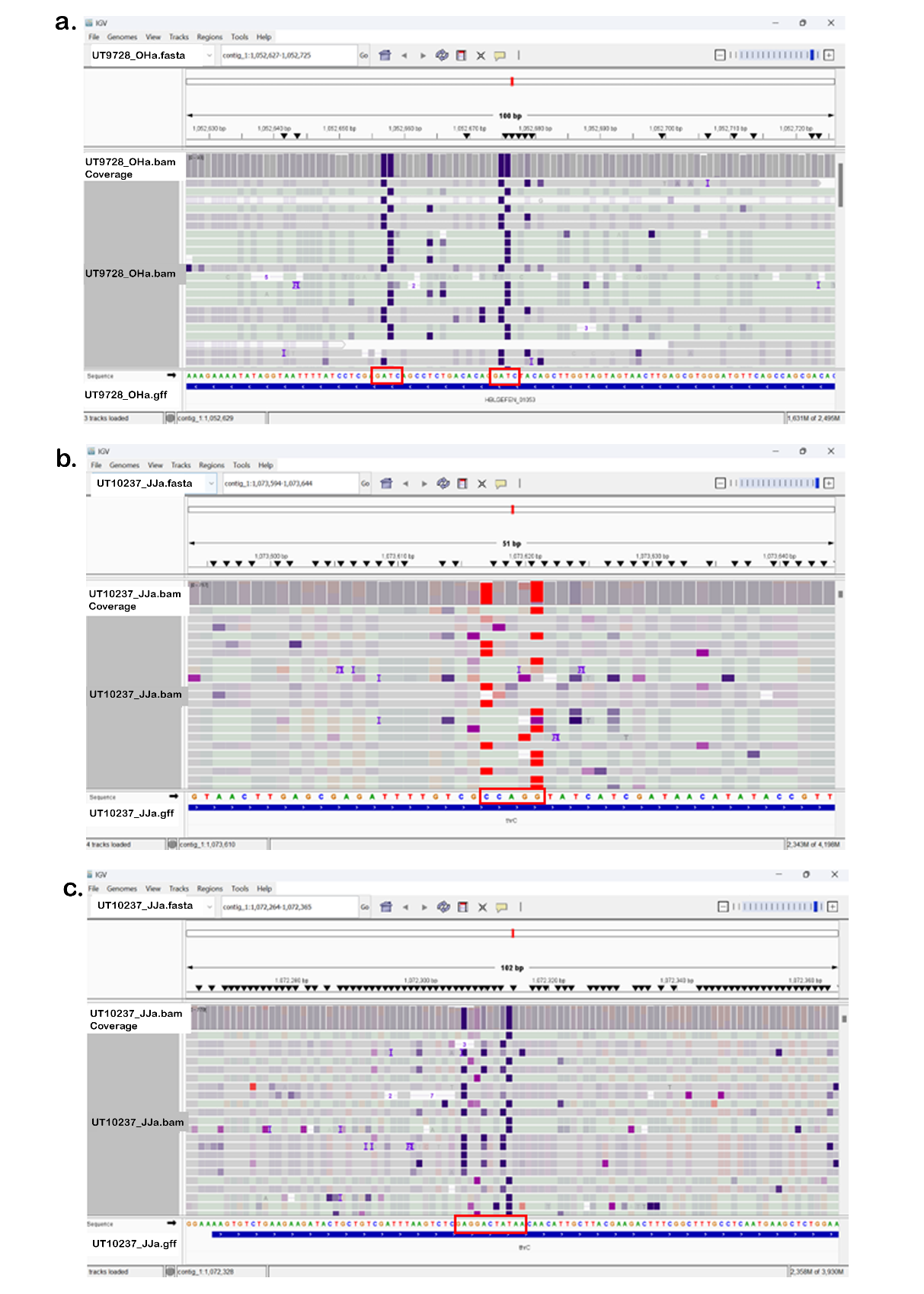

**Figure S3: IGV snapshots illustrating methylation at identified motif sites in ONT-only assemblies.** IGV tracks displaying base modification signals at key methylation motifs detected by MicrobeMod. (a) Methylation (purple) at the GATC motif in strain UT9728. (b) Methylation (red) at the CCWGG motif in strain UT10237. (c) Methylation (purple) at the GAGNNNNNTAA motif in strain UT10237.

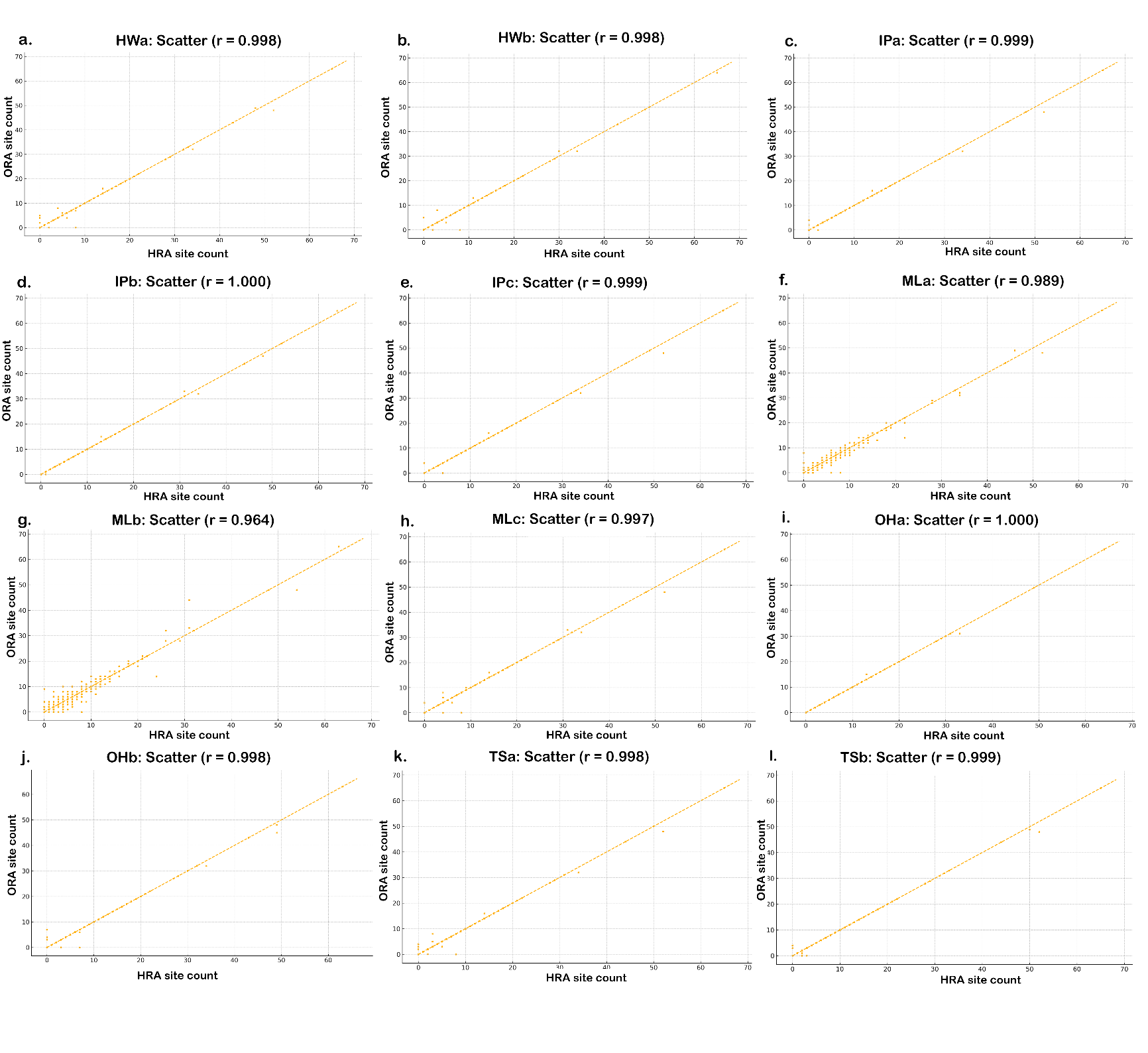

**Figure S4. UT9728 replicates show near-perfect linear relationship.** Individual scatter plots for all replicates of UT9728 HRA- and ORA- based methylation site calls. Dispersion is mostly limited to the MLb, and to some extend in the MLa replicates.

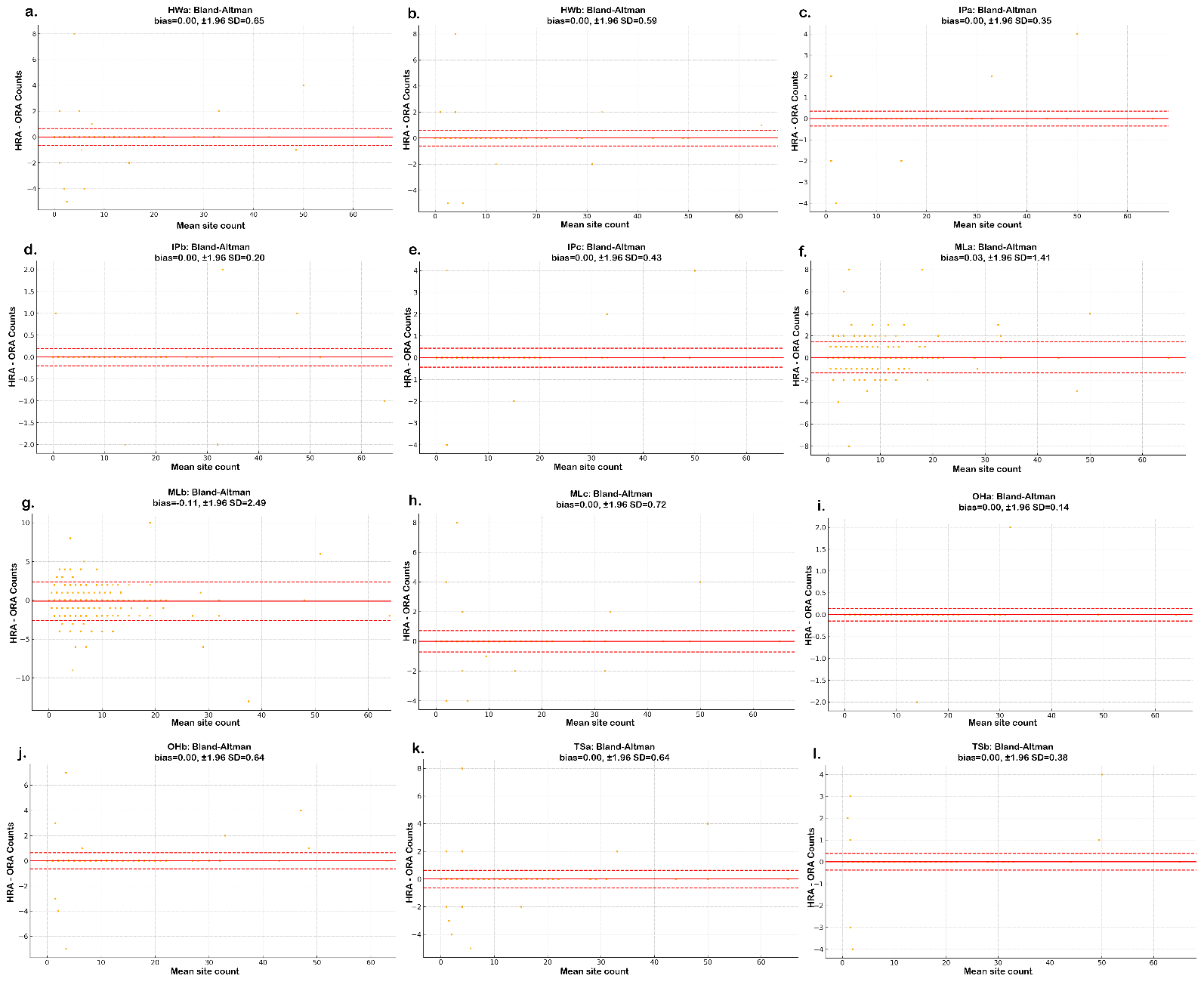

**Figure S5: UT9728 replicates indicate minimal bias.** Bland-Altman plots for each replicate of UT9728 HRA- and ORA- based methylation site calls. Bias is evident in MLb, and to some extent, in MLa replicate.

**Supplementary Tables**

| **Assembly** | **Motif*** |
| --- | --- |
| UT9728_HWa | 99.8% |
| UT9728_HWb | 99.8% |
| UT9728_IPa | 99.9% |
| UT9728_IPb | 99.9% |
| UT9728_IPc | 99.8% |
| UT9728_MLa | 99.9% |
| UT9728_MLb | 99.9% |
| UT9728_MLc | 99.9% |
| UT9728_OHa | 99.9% |
| UT9728_OHb | 99.9% |
| UT9728_TSa | 99.9% |
| UT9728_TSb | 99.9% |
| UT10237_JJa | 100.3%/100% |
| UT01237_JJb | 101.6%/100% |
| UT10237_TSc | 99.9%/99.1% |

**Table S1: MMC Analysis for all replicates.** Table detailing the motif methylation concordance for all replicates. Strain UT9728 motif is GATC and strain UT10237 motifs are GAGNNNNNTAA/CCWGG.

**a.**

| Replicate | Total sites | Mean percent modified | Strand + | | Strand - |
| --- | --- | --- | --- | --- | --- |
| UT9728_Hwa | 8486 | 0.855 | 4252 | 4234 | |
| UT9728_HWb | 8296 | 0.855 | 4160 | 4136 | |
| UT9728_IPa | 8487 | 0.854 | 4250 | 4237 | |
| UT9728_IPb | 8367 | 0.849 | 4191 | 4176 | |
| UT9728_IPc | 8436 | 0.852 | 4222 | 4214 | |
| UT9728_MLa | 8399 | 0.846 | 4217 | 4182 | |
| UT9728_MLb | 8430 | 0.852 | 4221 | 4209 | |
| UT9728_MLc | 8462 | 0.854 | 4236 | 4226 | |
| UT9728_OHa | 8281 | 0.846 | 4125 | 4156 | |
| UT9728_OHb | 8271 | 0.847 | 4137 | 4134 | |
| UT9728_TSa | 8355 | 0.845 | 4175 | 4180 | |
| UT9728_TSb | 8440 | 0.848 | 4225 | 4215 | |
| UT10237_JJa | 3839 | 0.794 | 1976 | 1863 | |
| UT10237_JJb | 3766 | 0.795 | 1936 | 1803 | |
| UT10237_TSc | 3741 | 0.795 | 1939 | 1802 | |

**b.**

| Replicate | Total sites | Mean percent modified | Strand + | | Strand - |
| --- | --- | --- | --- | --- | --- |
| UT9728_Hwa | 8486 | 0.855 | 4253 | 4233 | |
| UT9728_HWb | 8297 | 0.855 | 4160 | 4137 | |
| UT9728_IPa | 8487 | 0.855 | 4250 | 4237 | |
| UT9728_IPb | 8368 | 0.849 | 4190 | 4178 | |
| UT9728_IPc | 8436 | 0.852 | 4222 | 4214 | |
| UT9728_MLa | 8399 | 0.846 | 4217 | 4182 | |
| UT9728_MLb | 8430 | 0.852 | 4221 | 4209 | |
| UT9728_MLc | 8465 | 0.854 | 4237 | 4228 | |
| UT9728_OHa | 8281 | 0.846 | 4125 | 4156 | |
| UT9728_OHb | 8267 | 0.848 | 4136 | 4131 | |
| UT9728_TSa | 8355 | 0.845 | 4175 | 4180 | |
| UT9728_TSb | 8438 | 0.849 | 4223 | 4215 | |
| UT10237_JJa | 3860 | 0.794 | 1985 | 1875 | |
| UT10237_JJb | 3790 | 0.795 | 1973 | 1817 | |
| UT10237_TSc | 3744 | 0.795 | 1941 | 1803 | |

**Table S2: Total methylation site calls versus strand distribution.** Table detailing the total methylation site calls from MicrobeMod, the mean percent modification, and strand counts for all replicates of hybrid (a) and ONT (b) assemblies.

**a.**

| **Replicate** | **TP** | **FP** | **FN** | **TN** | **Sensitivity** | **Specificity** | **Precision** | **F1** |
| --- | --- | --- | --- | --- | --- | --- | --- | --- |
| HWa | 1452 | 2 | 0 | 3 | 1 | 0.6 | 0.999 | 0.999 |
| HWb | 1451 | 2 | 0 | 4 | 1 | 0.667 | 0.999 | 0.999 |
| IPa | 1454 | 1 | 0 | 2 | 1 | 0.667 | 0.999 | 1 |
| IPb | 1449 | 1 | 0 | 7 | 1 | 0.875 | 0.999 | 1 |
| IPc | 1450 | 1 | 0 | 6 | 1 | 0.857 | 0.999 | 1 |
| MLa | 1441 | 4 | 11 | 1 | 0.992 | 0.2 | 0.997 | 0.995 |
| MLb | 1399 | 5 | 51 | 2 | 0.965 | 0.286 | 0.996 | 0.98 |
| MLc | 1450 | 3 | 0 | 4 | 1 | 0.571 | 0.998 | 0.999 |
| OHa | 1452 | 0 | 0 | 5 | 1 | 1 | 1 | 1 |
| OHb | 1448 | 2 | 0 | 7 | 1 | 0.778 | 0.999 | 0.999 |
| TSa | 1449 | 2 | 0 | 6 | 1 | 0.75 | 0.999 | 0.999 |
| TSb | 1452 | 2 | 0 | 3 | 1 | 0.6 | 0.999 | 0.999 |

**b.**

| **Replicate** | **TP** | **FP** | **FN** | **TN** | **Sensitivity** | **Specificity** | **Precision** | **F1** |
| --- | --- | --- | --- | --- | --- | --- | --- | --- |
| JJa | 848 | 5 | 0 | 14 | 1 | 0.736842 | 0.994138 | 0.997061 |
| JJb | 840 | 4 | 1 | 22 | 0.998811 | 0.846154 | 0.995261 | 0.997033 |
| TSc | 827 | 4 | 0 | 36 | 1 | 0.9 | 0.995187 | 0.997587 |

**c.**

| **Replicate** | **TP** | **FP** | **FN** | **TN** | **Sensitivity** | **Specificity** | **Precision** | **F1** |
| --- | --- | --- | --- | --- | --- | --- | --- | --- |
| JJa | 379 | 0 | 0 | 43 | 1 | 1 | 1 | 1 |
| JJb | 360 | 0 | 1 | 61 | 0.99723 | 1 | 1 | 0.998613 |
| TSc | 365 | 0 | 0 | 57 | 1 | 1 | 1 | 1 |

**Table S3: Confusion matrix per motif.** Confusion matrices for UT9728 replicates (a), UT10237 replicates for CCWGG motif (b), and UT10237 replicates for GAGNNNNNTAA motif (c).
